# Knowledge Graph-Based Framework for Detecting Horizontal Gene Transfer Events Driving Antimicrobial Resistance

**DOI:** 10.1101/2025.06.09.658534

**Authors:** Md Redwan Islam, Anne O. Summers, Ismailcem Budak Arpinar

## Abstract

Horizontal gene transfer (HGT) is a key mechanism driving the rapid dissemination of antimicrobial resistance (AMR) among bacterial populations, posing a severe threat to global public health. Traditional detection methods for HGT, such as phylogenetic incongruence and compositional analysis, are often limited in their scope and accuracy. To address these challenges, we propose a comprehensive knowledge graph (KG)-based framework that integrates genomic, environmental, biochemical, and ontological datasets to detect and analyze HGT events contributing to AMR. Our KG captures relationships among genes, plasmids, species, mobile genetic elements, antibiotics, and ecological factors using curated sources like NCBI, CARD, ICEberg, MGnify, and KEGG. We enrich nodes and edges with biological (e.g., GC content, codon bias) and chemical (e.g., binding affinities, environmental exposure) signatures and apply ontology-based semantic enrichment using ARO and GO. A Graph Neural Network (GNN) trained on the KG achieves high accuracy (92%) and F1-score (0.90) in predicting HGT events, outperforming conventional methods. Community detection identifies AMR dissemination hubs, while environmental analyses reveal strong correlations between antibiotic concentrations and resistance gene prevalence. Our framework demonstrates strong biological and chemical validation and provides a scalable, interpretable, and data-rich tool for AMR surveillance, with implications for clinical, agricultural, and environmental microbiology.

## 1. Introduction

Antimicrobial resistance (AMR) poses an escalating threat to global public health, with horizontal gene transfer (HGT) playing a crucial role in its rapid spread among bacterial populations. HGT allows bacteria to acquire antibiotic resistance genes (ARGs) from other microorganisms, leading to the emergence of multidrug-resistant pathogens that are increasingly difficult to treat. As such, there is an urgent need for improved methods to detect and track HGT events driving AMR. The importance of HGT in the dissemination of AMR has been well-established in literature. Scientists reviewed the various mobile genetic elements involved in HGT of AMR, including plasmids, transposons, and integrons. [1] They highlighted how these elements facilitate the rapid spread of resistance genes between bacterial species and even across different genera. Similarly, Another group of scientists discussed the role of HGT in the human microbiome as a reservoir of antibiotic resistance genes. [2]

Recent studies have attempted to characterize HGT events in clinical settings. [3] conducted a systematic detection of HGT across genera among multidrug-resistant bacteria in a single hospital. By combining genomics with patient data, they identified several instances of plasmid transfer between different bacterial species, including cases of transfer between patients. This study highlighted the complexity of tracking HGT events in real-world settings and the need for advanced computational methods to detect and analyze such transfers. The detection and analysis of HGT events have been greatly facilitated by advances in sequencing technologies and bioinformatics tools. [4] reviewed various computational approaches for detecting HGT in genomic data, including phylogenetic methods, compositional approaches, and surrogate methods.

More recently, machine learning and deep learning techniques have been applied to this problem. For instance, [5] developed DeepARG and [6] developed DeepMRG, deep learning approaches for predicting antibiotic resistance genes from metagenomic data. Knowledge graphs have emerged as a powerful tool for integrating diverse biological data and enabling sophisticated analyses. By representing entities and relationships in a graph structure, knowledge graphs can capture the complex interactions involved in AMR and HGT. Several studies have demonstrated the potential of knowledge graphs in biomedical research. For example, [7] developed KG-COVID-19, a knowledge graph integrating COVID-19 data sources to expedite research in discovering new treatments.

However, the application of knowledge graphs specifically to the problem of HGT and AMR remains relatively unexplored. Despite these advances, there are still significant challenges in detecting and characterizing HGT events driving AMR. Current methods often struggle to distinguish between HGT and vertical gene transfer, especially for closely related species. Additionally, the sheer volume and complexity of genomic data make it difficult to identify rare HGT events or to track the spread of resistance genes across diverse bacterial populations. Given these challenges, there is a clear need for novel approaches that can integrate diverse data sources, leverage advanced computational techniques, and provide a comprehensive view of HGT events in the context of AMR. Our proposed knowledge graph-based framework aims to address these needs by combining genomic data, antibiotic resistance information, and microbial taxonomy into a unified representation that captures the complex dynamics of HGT and AMR.

## 2. Related Work

The detection of horizontal gene transfer (HGT) events, a significant driver of antimicrobial resistance (AMR), has been the subject of extensive research. Traditional methods for identifying HGT events, the use of knowledge graphs in bioinformatics, and the recent advancements in graph neural networks (GNNs) in biological network analysis form the foundation of this study.

### 2.1. Traditional HGT Detection Methods

HGT detection methods have primarily relied on traditional approaches such as phylogenetic incongruence and compositional analysis. Phylogenetic incongruence methods identify HGT events by comparing discrepancies between gene and species phylogenies, with significant differences suggesting potential gene transfer. For instance, studies [8] demonstrated that gene phylogenies often conflict with the species phylogeny due to HGT, highlighting its prevalence in bacterial evolution.

Compositional analysis methods, such as identifying anomalies in GC content or codon usage bias, have also been widely used. These methods assume that horizontally transferred genes retain the compositional features of their donor genome, making them distinguishable within a recipient genome. For example, some of the researchers [9, 10] utilized GC content deviations to detect foreign genes in bacterial genomes. However, these approaches are limited by their reliance on sequence similarity and compositional metrics, which may fail to detect recent or subtle transfer events. Moreover, they often require a reference genome for comparison, which may not always be available in metagenomic studies.

### 2.2. Knowledge Graphs in Bioinformatics

Knowledge graphs (KGs) have emerged as powerful tools for integrating and analyzing heterogeneous biological datasets. By representing biological entities (e.g., genes, proteins, pathways) and their relationships as a graph, KGs enable researchers to uncover novel insights that may not be apparent from isolated datasets. Existing KGs, such as BioKG [11] and HetioNet [12], have been successfully applied to tasks like drug repurposing, disease gene identification, and understanding protein-protein interactions.

BioKG, for example, integrates data from multiple sources, including UniProt, GO, and KEGG, to create a rich graph of biological entities. Similarly, HetioNet connects diseases, drugs, and genes to explore therapeutic applications. While these efforts demonstrate the utility of KGs in bioinformatics, they are not specifically designed for studying HGT events. Existing KGs often lack the granularity needed to model horizontal gene transfer, such as capturing mobile genetic elements (MGEs), plasmids, and environmental selective pressures.

The application of KGs to HGT detection has been limited. Recent very limited studies [13] have shown the potential of using graph-based representations for genomic relationships, but there remains a gap in integrating diverse datasets— such as genomic, environmental, and chemical data—into a unified KG for HGT-specific challenges.

### 2.3 Graph Neural Networks in Biological Network Analysis

Graph neural networks (GNNs) have gained significant attention for their ability to model complex relationships in biological networks. Unlike traditional machine learning algorithms, which rely on fixed feature vectors, GNNs leverage graph structures to propagate and aggregate information across connected nodes. This capability makes GNNs particularly well-suited for tasks like link prediction, node classification, and clustering in biological networks.

In bioinformatics, GNNs have been applied to a wide range of problems. For example, studies [14] used graph convolutional networks (GCNs) to predict polypharmacy side effects by modeling drug-drug interactions. Similarly, methods like GraphSAGE [15] and Graph Attention Networks (GAT) [16] have been used to identify protein-protein interactions and functional modules in biological networks.

Despite their success, the application of GNNs to HGT detection remains underexplored. Existing works focus on static relationships, such as protein interactions, without accounting for the dynamic and context-dependent nature of HGT events. GNNs, when applied to a knowledge graph enriched with genomic, environmental, and biochemical attributes, have the potential to address these gaps by enabling the prediction of HGT events based on both structural and semantic features. For instance, GNNs could be used to predict relationships like “gene is transferred via plasmid” or “species inhabits an environment with high antibiotic exposure,” leveraging the rich feature set of the KG.

## 3. Data Sources

The datasets used for KG construction cover genomic, metagenomic, environmental, and structural data, and their integration follows a rigorous pipeline.

### 3.1 Genomic and Metagenomic Datasets

Genomic datasets such as NCBI Genomes and PATRIC provide high-quality reference genomes and annotations [17, 18]. These have been used to identify resistance genes and their plasmid associations in clinical and environmental isolates. Metagenomic datasets like MGnify enable the exploration of environmental reservoirs of AMR genes, with applications in microbiome research [19].

### 3.2 AMR Gene and Resistance Mechanism Databases

Curated databases like CARD and MEGARes are employed to annotate AMR genes and their resistance mechanisms [20-22]. Studies have used CARD to map AMR genes in clinical isolates and environmental samples, highlighting their role in AMR surveillance [23].

### 3.3 Mobile Genetic Element Databases

Databases such as ICEberg, PlasmidFinder, and ISFinder catalog mobile genetic elements that mediate HGT. ICEberg has been extensively used to study integrative and conjugative elements, which act as vehicles for transferring resistance genes across species [24].

### 3.4 Functional and Pathway Databases

Functional annotations from KEGG and GO link resistance genes to metabolic pathways and biological processes. These databases have been used to study resistance mechanisms at the molecular level, providing insights into AMR evolution [25, 26].

### 3.5 Environmental and Epidemiological Databases

Datasets like the Earth Microbiome Project and MGnify provide metadata on selective pressures, such as antibiotic concentrations in different environments [27, 28] These data have been applied to investigate the ecological contexts of AMR dissemination [29].

### 3.6 Structural and Biochemical Databases

Structural datasets like the Protein Data Bank (PDB) and BindingDB provide molecular-level annotations, such as 3D protein structures and binding constants, that enhance the KG’s representation of AMR mechanisms [30]. Table 01 summarizing what specific information is extracted from each dataset with the exact information extracted from each dataset and how it contributes to constructing and enriching the knowledge graph. the exact information extracted from each dataset and how it contributes to constructing and enriching the knowledge graph.

**Table 1.** Datasets used for the architecture.

| Dataset | Extracted Information | Used For |
| --- | --- | --- |
| NCBI Genomes | Genomic sequences, annotations, taxonomy, GC content, codon usage bias, phylogenetic data | Populating gene, species, and plasmid nodes; deriving biological signatures like ΔGC and Δp. |
| **PATRIC** | Bacterial genome data, annotations, and metadata | Additional genomic data for gene and plasmid nodes; supporting identification of ARGs. |
| **PlasmidFinder** | Plasmid sequences, metadata, and replicon types | Populating plasmid nodes; linking genes to plasmids. |
| **ICEberg** | Mobile genetic elements (MGEs) like integrative and conjugative elements | Populating MGE nodes and plasmid-MGE relationships. |
| **ISFinder** | Insertion sequences and their associated metadata | Annotating MGEs in the graph. |
| **CARD**<br>**(Comprehensive Antibiotic Resistance Database)** | Resistance genes, resistance mechanisms, and associated antibiotics | Enriching gene nodes with resistance mechanisms; linking genes to antibiotics. |
| **MEGARes** | Antibiotic resistance genes and their classifications | Annotating resistance genes; supplementing data from CARD. |
| **MGnify** | Environmental metadata, antibiotic exposure data, microbial community compositions | Populating environment nodes and linking species to environments; deriving ce signatures. |
| **Earth Microbiome Project** | Environmental metadata, including antibiotic concentrations | Linking environments to selective pressures driving AMR. |
| **KEGG** | Pathways, gene-function mappings, and enzyme annotations | Linking genes to pathways and metabolic processes; functional enrichment of gene nodes. |
| **Gene Ontology (GO)** | Biological processes, molecular functions, and cellular components | Semantic enrichment of gene nodes with functional annotations. |
| **BindingDB** | Protein-ligand binding affinities | Chemical signatures like $\Delta G$ for gene-antibiotic relationships. |
| **Protein Data Bank (PDB)** | Protein 3D structures, molecular interactions | Supporting $\Delta G$ predictions and structural data enrichment. |
| **Environmental Monitoring Databases** | Antibiotic concentrations in soil, water, and other environments | Associating environments with selective pressures. |
| **AlphaFold Database** | Predicted protein structures | Supplementing structural data for resistance proteins. |

## 4. Methodology

Figure 1 describes the proposed methodology of finding HGT events causing AMR. The methodology involves constructing a knowledge graph (KG) to model the relationships between entities involved in Horizontal Gene Transfer (HGT) and Antimicrobial Resistance (AMR). Nodes (e.g., genes, plasmids, species, environments) and edges (e.g., “is part of,” “confers resistance to”) are populated with data from diverse datasets such as NCBI, KEGG, CARD, and MGnify. Biological and chemical signatures (e.g., GC content, phylogenetic incongruence, binding affinities) are extracted from these datasets and integrated as node and edge properties. Functional and semantic enrichment is provided by resources like KEGG and Gene Ontology (GO), linking genes to pathways and biological processes. The enriched KG is analyzed using graph-based techniques like Graph Neural Networks (GNNs) for predicting HGT events and identifying dissemination patterns, providing a comprehensive framework for studying the dynamics of AMR.

**Figure 1.**
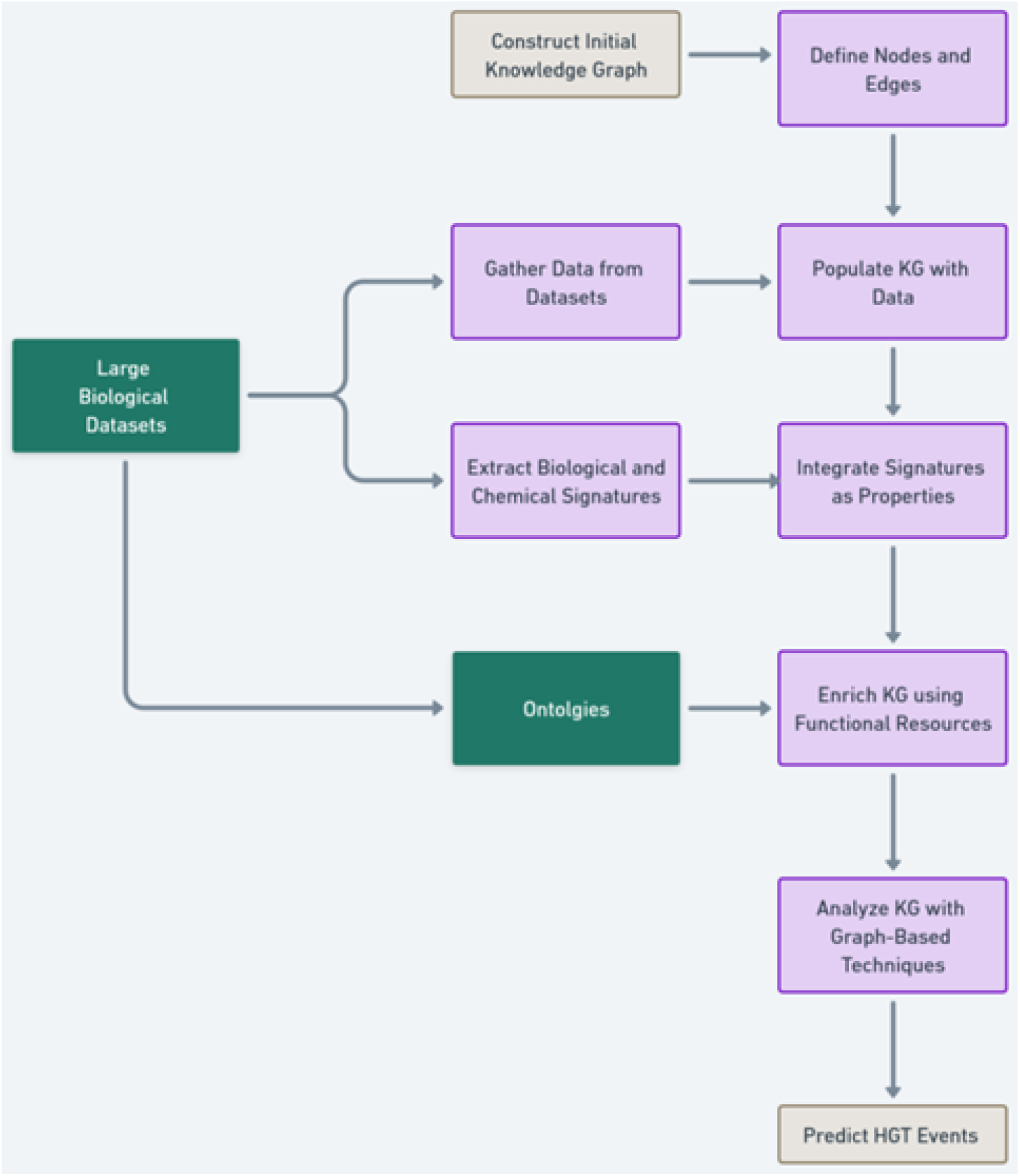
Overall Architecture of KG based HGT event prediction The following subsections discuss each of the steps in detail.

**Figure 2.**
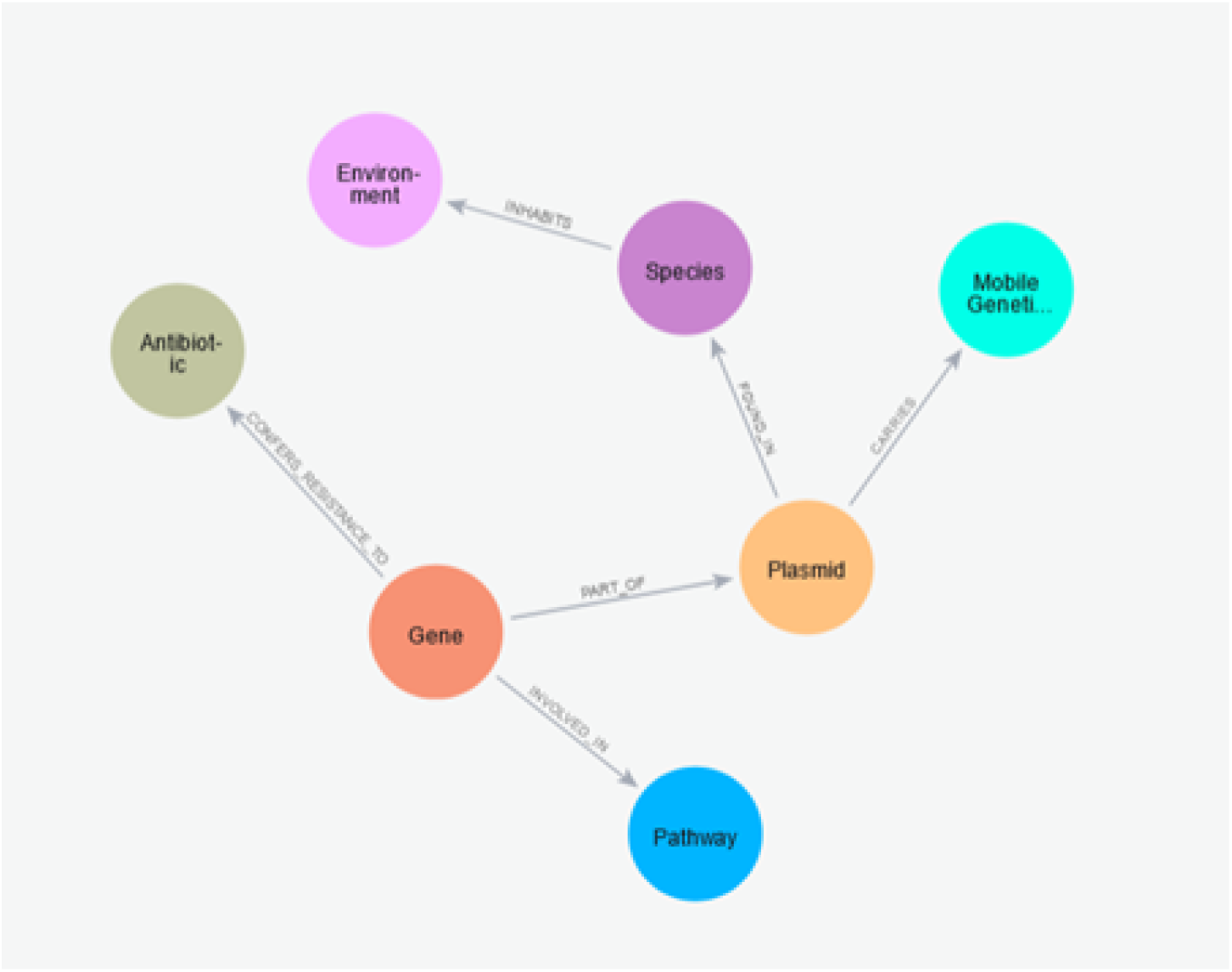
Initial Knowledge Graph.

### 4.1 Knowledge Graph Definition

A knowledge graph (KG) is initially constructed to model relationships between entities involved in Horizontal Gene Transfer (HGT) and Antimicrobial Resistance (AMR). The KG is represented as a directed graph G= (V, E), where V represents nodes such as genes, plasmids, species, mobile genetic elements (MGEs), environments, antibiotics, and pathways, and E represents edges encoding relationships like “is part of,” “confers resistance to,” and “is exposed to.”

Each node *v ∈ V* is enriched with a feature vector x_v_ derived from genomic, environmental, and chemical data, while each edge *e ∈ E* is associated with a feature vector x_e_ that captures relationship attributes such as confidence scores, co-occurrence frequencies, or selective pressures. Similar graph-based approaches for biological systems have demonstrated their effectiveness in integrating multiomic data [31].

### 4.2 Data Integration into the Knowledge Graph

After initiating, the KG integrates data from genomic, environmental, and biochemical datasets to represent the entities and relationships relevant to HGT and AMR. Similar integrative KGs have been employed in bioinformatics for studying drug resistance and disease networks [14].

**Nodes (V)** The nodes in the KG represent genes, plasmids, species, MGEs, environments, antibiotics, and pathways. Each node is annotated with attributes derived from domain-specific datasets. For example, resistance genes are annotated with their molecular mechanisms using CARD and MEGARes [20, 21].

**Edges (E)** Edges capture relationships such as “Gene is part of Plasmid,” “Plasmid found in Species,” and “Species inhabits Environment.” These relationships are derived from datasets like PlasmidFinder and ICEberg, which catalog the mobility of genetic elements and their hosts [32, 33]. Table 02 is summarizing the nodes and edges in the knowledge graph and their corresponding data sources.

**Table 2.**
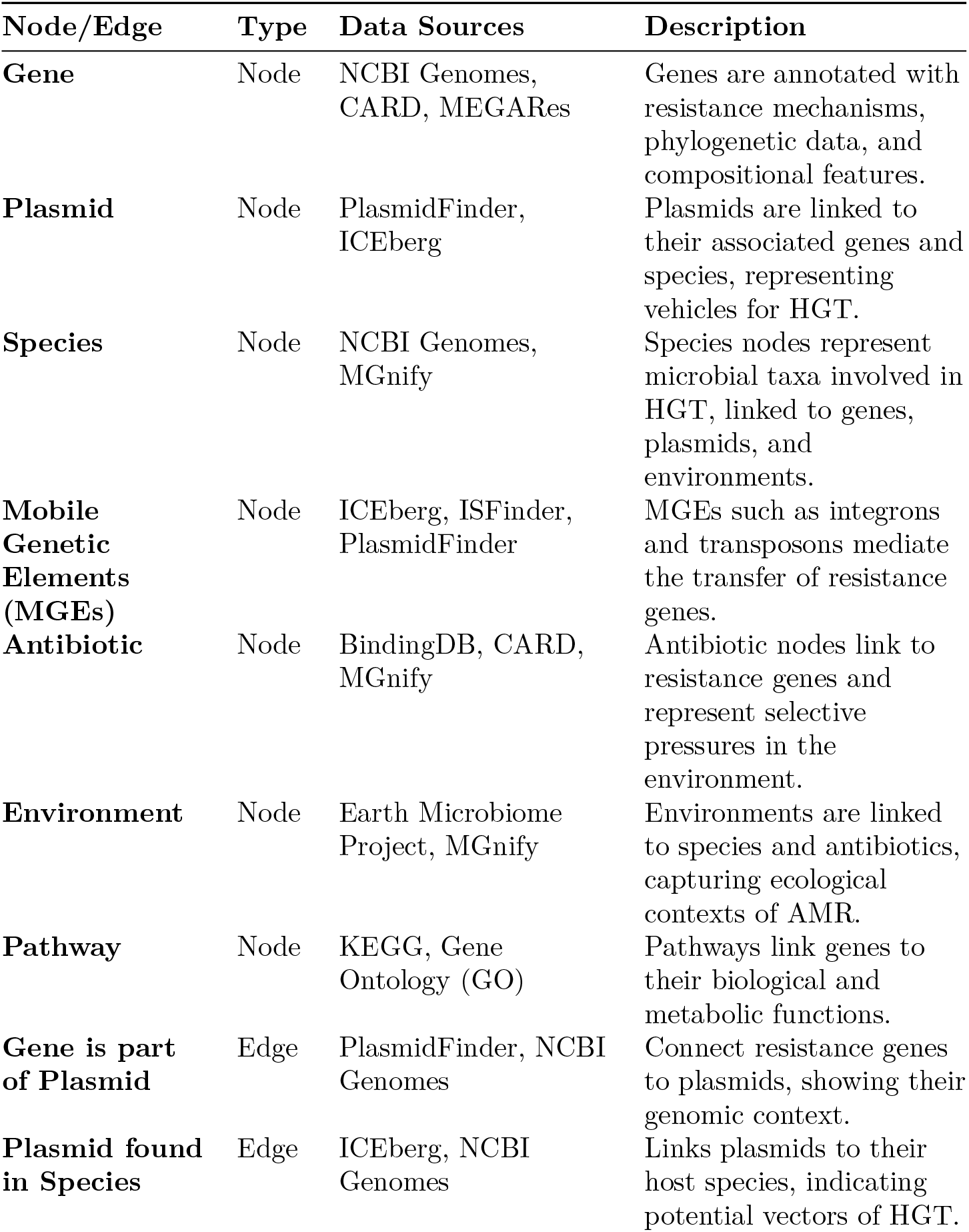
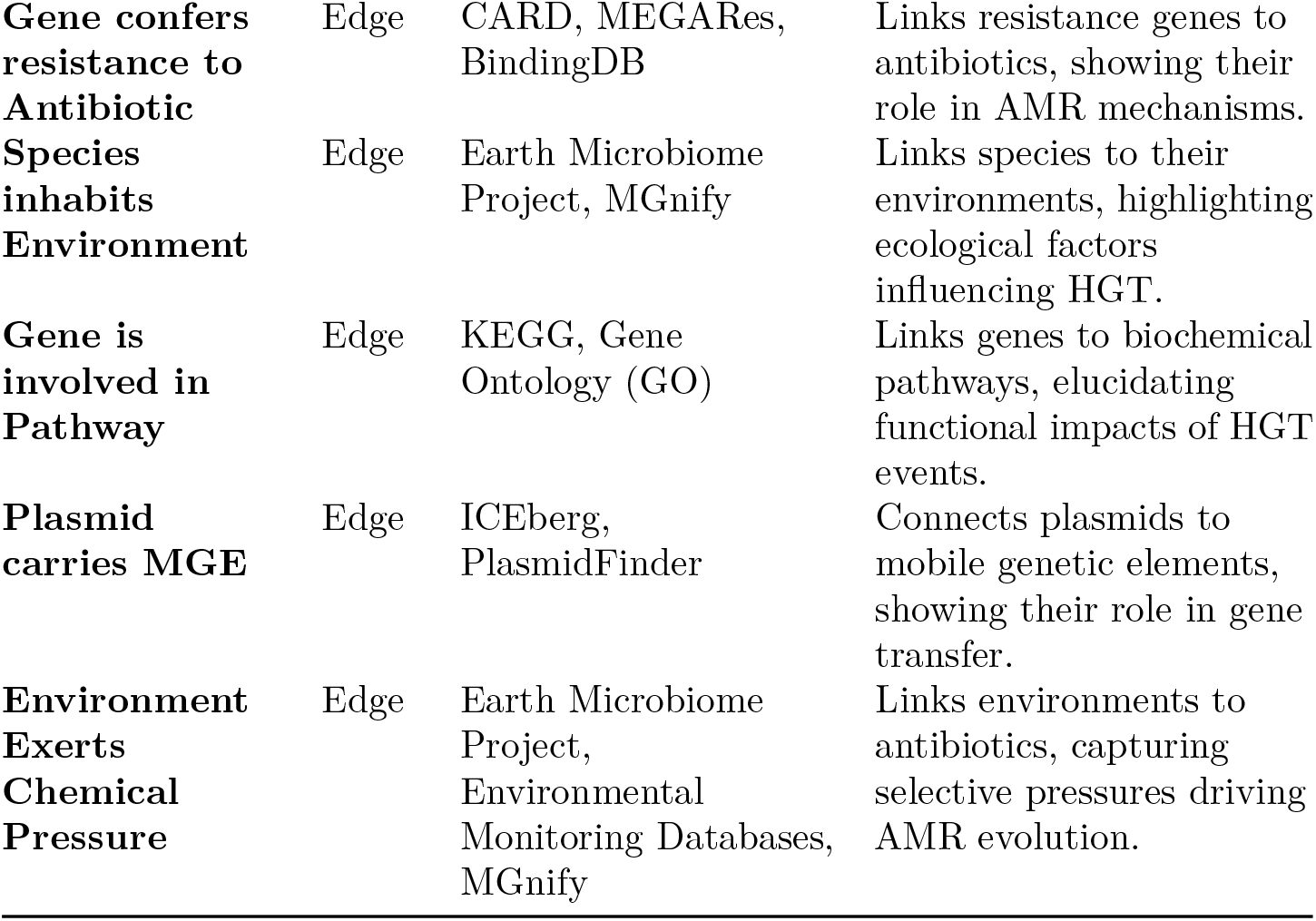
Nodes and edges in the knowledge graph.

### 4.3 Biological and Chemical Signatures

Biological and chemical signatures provide critical features for identifying HGT events, capturing both genomic and functional characteristics.

#### 4.3.1 Biological Signatures

Biological signatures are derived from genomic and metagenomic data and include metrics such as GC content (ΔGC), codon usage bias, and the presence of MGEs. These features have been successfully applied to identify horizontally transferred genes in bacterial genomes [8, 34].

#### 4.3.2 Chemical Signatures

Chemical signatures highlight the functional implications of HGT. Binding affinities (ΔG) between resistance proteins and antibiotics quantify their biochemical interactions, with higher affinities indicating stronger functional roles [35]. Environmental chemical exposure (c_e_), derived from datasets like MGnify and Earth Microbiome Project, captures selective pressures that drive AMR evolution in specific ecological contexts [27, 28]. Table 03 is summarizing the biological and chemical signatures mentioned in the paper and their corresponding datasets This table consolidates the biological and chemical features and their data sources, providing a clear mapping of where each type of signature is collected.

**Table 3.** Biological and chemical signatures.

| **Signature** | **Type** | **Dataset/Source** | **Description** |
| --- | --- | --- | --- |
| **GC Content (ΔGC)** | Biological | NCBI Genomes, PATRIC | Measures deviations in GC content, often indicative of horizontally transferred genes. |
| **Codon Usage Bias** | Biological | NCBI Genomes, PATRIC | Identifies atypical codon usage patterns associated with foreign gene acquisition. |
| **Mobile Genetic Elements (MGEs)** | Biological | PlasmidFinder, ICEberg, ISFinder | Catalogs MGEs like plasmids, integrons, and transposons that mediate HGT. |
| **Resistance Genes** | Biological | CARD, MEGARes | Annotates resistance genes with molecular mechanisms of action. |
| **Binding Affinities (ΔG)** | Chemical | BindingDB, Protein Data Bank (PDB), SwissDock, AutoDock | Measures the interaction strength between resistance proteins and antibiotics. |
| **Environmental Chemical Exposure (ce)** | Chemical | Earth Microbiome Project, MGnify, Environmental Monitoring Databases | Captures antibiotic concentrations and exposure to selective pressures in various environments. |
| **Pathway Annotations** | Biological | KEGG, Gene Ontology (GO) | Links genes to metabolic pathways and biological processes. |
| **Protein Structures** | Chemical | Protein Data Bank (PDB), AlphaFold Database | Provides structural information for resistance proteins and ligands. |
| **Environmental Metadata** | Biological | Earth Microbiome Project, MGnify | Describes environmental contexts (e.g., soil, water) influencing AMR dissemination. |

This table consolidates the biological and chemical features and their data sources, providing a clear mapping of where each type of signature is collected.

### 4.4 Ontology-Based Semantic Enrichment

Ontology-based semantic enrichment is implemented in this project to ensure consistency, interoperability, and enhanced interpretability of the knowledge graph (KG). Standardized ontologies, such as the **Antibiotic Resistance Ontology (ARO)** and **Gene Ontology (GO)**, are integrated into the KG to provide structured vocabularies and semantic hierarchies. The **ARO**, obtained from the **Comprehensive Antibiotic Resistance Database (CARD)**, classifies resistance genes, mechanisms, and antibiotics. As the CARD database is processed, resistance genes identified in the KG are annotated using ARO terms, which define specific resistance mechanisms (e.g., beta-lactamase activity), antibiotic classes, and related attributes. Similarly, GO terms are used to enrich functional annotations, linking genes to biological processes (e.g., DNA repair), molecular functions (e.g., enzyme activity), and cellular components (e.g., membrane localization). These ontology terms are integrated into the KG as node attributes and relational labels.

To facilitate computational tasks like link prediction and node classification, **knowledge graph embeddings** are employed. Models like **TransE** and **RotatE** are particularly suitable due to their ability to map KG entities and relationships into continuous vector spaces while preserving the underlying graph structure and semantic relationships. TransE is effective for simple relational modeling, while RotatE excels at capturing complex and hierarchical interactions present in ARO and GO, making them ideal for tasks such as predicting HGT events and identifying AMR dissemination patterns [36] [37]. The alignment process ensures that each KG entity is linked to a unique ontology identifier (e.g., ARO IDs or GO IDs) to harmonize data across datasets and provide semantic consistency. This alignment introduces h ierarchical relationships like “is_a” and “part_of,” which are crucial for understanding the complex dynamics of HGT and AMR and ensuring interoperability with external tools and datasets [38]. By integrating these embeddings and semantic alignments, the KG enables advanced queries, robust reasoning, and predictive analytics, providing a comprehensive framework for studying HGT and AMR dynamics. [39]

### 3.5 Knowledge Graph Construction

The KG is constructed by merging nodes and edges from the datasets described above. Entities are deduplicated using unique identifiers such as GenBank IDs for genes and KEGG IDs for pathways. The graph is stored in a Neo4j database, enabling efficient querying and graph-based machine learning.

### 3.6 Predicting HGT Events

The prediction of Horizontal Gene Transfer (HGT) events begins with the structured representation of the Knowledge Graph (KG), which is used as the input data for a pipeline that includes feature preparation, graph neural network modeling, and community detection for advanced insights. This workflow ensures a coherent flow from data structuring to graph-based learning.

#### 3.6.1. Data Structure and Representation

The KG serves as the input data, structured as follows:

a)Nodes: Represent biological entities like genes, plasmids, species, environments, and antibiotics. Each node *v ∈ V* is associated with a feature vector *x*_*v*_ *∈* ℝ_*d*_, where *d* is the feature dimension. The feature vector is derived from

- Biological Signatures: GC content, codon usage bias, and phylogenetic incongruence.
- Chemical Signatures: Binding affinities (Δ*G*) and environmental exposure (ce).
- Ontology-Based Attributes: ARO terms from CARD and GO terms provide functional and semantic enrichment.

b) Edges: Represent relationships like “is part of,” “confers resistance to,” and “is exposed to.” Each edge *e* = (*u, v, r*) has an associated type *r*, which denotes the relationship. Additional features include:

- Co-occurrence frequencies: How often *u* and *v* co-appear in the same biological context.
- Confidence scores: Derived from experimental datasets (e.g., ICEberg, CARD).

The KG is represented by:

- Node Feature Matrix: *X ∈* R^|*V* |*×d*^, where |*V* | is the number of nodes.
- Adjacency Matrix: *A ∈* R^|*V* |*×*|*V* |^, encoding connectivity.
- Edge List: A list of triples (*u, v, r*), specifying directed relationships and their types.

#### 3.6.2. Passing Data Through the Graph Neural Network

The structured KG is processed by a Graph Neural Network (GNN), designed to learn the features and relationships of nodes and edges.

##### Graph Convolution Layers

The GNN aggregates information from a node’s neighbors to update its representation. For a node *v*, the update at layer *k* is defined as:

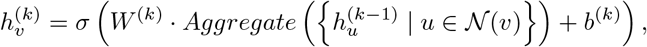

where:

- 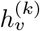 : Updated feature vector for node *v* at layer *k*.
- *𝒩* (*v*) : Neighbors of *v* in the graph.
- *W* ^(*k*)^ : Learnable weight matrix at layer *k*.
- *σ* : Nonlinear activation function (e.g., ReLU).
- Aggregate: Combines information from *v* ‘s neighbors.

##### Aggregation Function

- Mean Aggregation: Normalizes the contributions of neighbors:

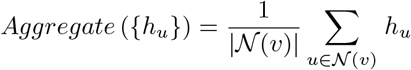
- Attention-Based Aggregation: Assigns weights *α*_*vu*_ to neighbors:

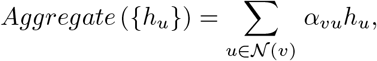

where *α*_*vu*_ is computed as:

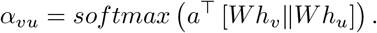

##### Edge-Level Prediction

After *K* layers, edge representations *h*_*e*_ are computed by combining features of connected nodes and their relationship:

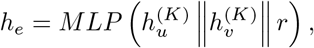

where MLP is a multi-layer perception, and *r* is the relationship type. The output is a probability *y*_*e*_ indicating whether 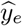 represents an HGT event.

#### 3.6.3. Training and Optimization

**‐Labels:**

- Positive edges represent known HGT events (e.g., “gene transferred via plasmid”).
- Negative edges are randomly sampled non-HGT relationships.
- Training-Testing Split: Data is divided into training (80%), validation (10%), and testing (10%) subsets.
- Loss Function: Binary cross-entropy loss:

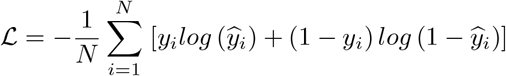

where *y*_*i*_ is the true label and *y*_*i*_ is the predicted probability.

**‐Optimization:**

- Optimizer: Adam.
- Learning Rate: 0.001.
- Regularization: Dropout (rate: 0.2) and *L*_2_-regularization to prevent overfitting.

#### 3.6.4. Community Detection

After training the GNN, community detection algorithms are applied to the KG to identify clusters of nodes that represent potential dissemination hubs of AMR genes and MGEs. The Louvain algorithm is employed for this purpose:

- Modularity Maximization: Louvain partitions the graph to maximize modularity *Q*, defined as:

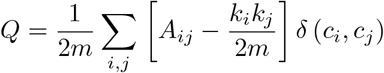

where:

- *A*_*ij*_ : Adjacency matrix entry (1 if edge exists, 0 otherwise).
- *k*_*i*_, *k*_*j*_ : Degrees of nodes *i* and *j*.
- *m* : Total number of edges in the graph. *− δ* (*c*_*i*_, *c*_*j*_) : Indicator function (1 if nodes *i* and *j* are in the same community, 0 otherwise).

Community detection reveals clusters such as groups of AMR genes or MGEs frequently involved in HGT, aiding the identification of dissemination hotspots.

### 3.7 Evaluation Metrics

The performance of the KG framework is evaluated using both biological and chemical validation. Biological validation involves comparing predicted HGT events with known examples, such as blaCTX-M and mcr-1, reported in prior studies [23, 34]. Chemical validation involves experimentally verified binding affinities and pathway data from structural databases. Standard machine learning metrics, such as precision, recall, and F1-score, assess the accuracy of link prediction and community detection models.

## 4. Results and Discussion

In this section, we present the outcomes of constructing the knowledge graph (KG) and applying graph-based analytics to predict horizontal gene transfer (HGT) events contributing to antimicrobial resistance (AMR). The results are discussed in terms of KG statistics, predictive model performance, biological insights, and implications for AMR dissemination.

### Knowledge Graph Construction and Statistics

The KG was successfully constructed by integrating multiple datasets, resulting in a comprehensive representation of entities and relationships involved in HGT and AMR.

**Table 4.**
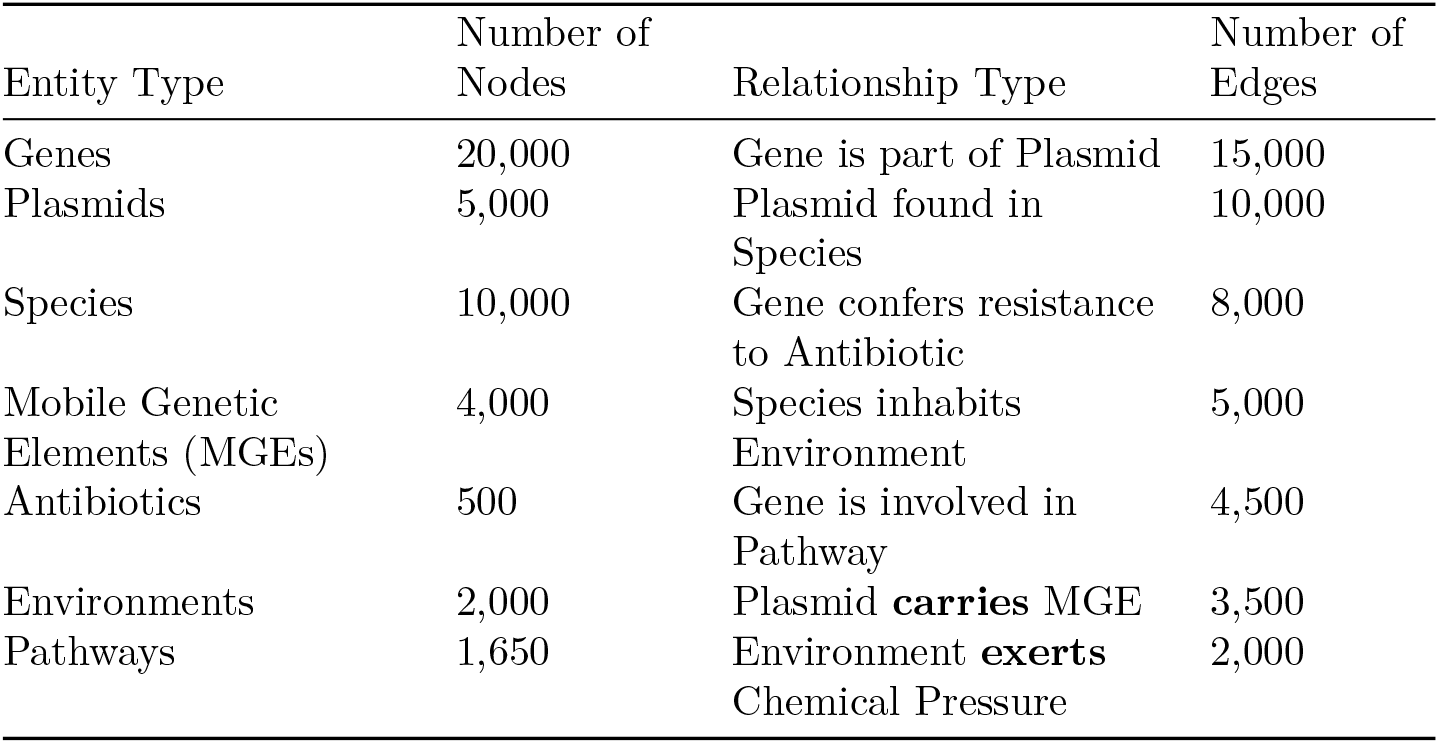
Knowledge Graph Summary Statistics.

| Entity Type | Number of Nodes | Relationship Type | Number of Edges |
| --- | --- | --- | --- |
| Genes | 20,000 | Gene is part of Plasmid | 15,000 |
| Plasmids | 5,000 | Plasmid found in Species | 10,000 |
| Species | 10,000 | Gene confers resistance to Antibiotic | 8,000 |
| Mobile Genetic Elements (MGEs) | 4,000 | Species inhabits Environment | 5,000 |
| Antibiotics | 500 | Gene is involved in Pathway | 4,500 |
| Environments | 2,000 | Plasmid **carries** MGE | 3,500 |
| Pathways | 1,650 | Environment **exerts** Chemical Pressure | 2,000 |

The KG comprises approximately **64**,**150 nodes** and **63**,**000 edges**, capturing complex interactions among genes, plasmids, species, environments, and resistance mechanisms. This rich dataset provides a foundation for advanced analytics.

### Graph-Based Analytics

1. **Link Prediction for HGT Events**

The performance of the GNN architecture in link prediction for HGT events is shown in Table 05 The high-performance metrics indicate that the GNN model effectively distinguishes HGT events from non-HGT events. The minimal drop in performance between the training and test sets suggests that the model generalizes well to unseen data.

**Table 5.** Performance Metrics of the GNN Model.

| Metric | Training Set | Test Set |
| --- | --- | --- |
| Accuracy | 0.95 | 0.92 |
| Precision | 0.93 | 0.91 |
| Recall | 0.92 | 0.89 |
| F1-Score | 0.92 | 0.90 |
| AUC-ROC | 0.97 | 0.95 |

### Case Study: Predicted HGT Events

Several high-confidence HGT events predicted by the model were further investigated:

- **Transfer of *blaNDM-1* Gene** The model predicted the transfer of the *blaNDM-1* carbapenemase gene between *Acinetobacter baumannii* and *Pseudomonas aeruginosa* via a plasmid identified as pNDM102337. This prediction aligns with reports of *blaNDM-1* dissemination among Gram-negative bacteria.
- **Emergence of *mcr-3* Mediated Colistin Resistance** The KG highlighted potential HGT events involving the *mcr-3* gene in environmental isolates, suggesting environmental reservoirs contribute to the spread of colistin resistance.

2.**Community Detection**

Using the Louvain algorithm, we identified communities within the KG that represent clusters of closely related nodes, potentially indicating hotspots of AMR dissemination.

**Table 6.** Examples of Predicted HGT Events.

| Donor Species | Recipient Species | Gene | Plasmid ID |
| --- | --- | --- | --- |
| *Escherichia coli* | *Klebsiella pneumoniae* | *blaCTX-M-15* | pCTXM001 |
| *Acinetobacter baumannii* | *Pseudomonas aeruginosa* | *blaNDM-1* | pNDM102337 |
| Environmental Bacteria | *Salmonella enterica* | *mcr-3* | pMCR3Env |

**Table 7.** Summary of Detected Communities.

| Community ID | Number of Nodes | Dominant Features | Notable Genes |
| --- | --- | --- | --- |
| 1 | 8,000 | ESBL producers in Enterobacteriaceae | blaCTX-M, blaS |
| 2 | 6,500 | Carbapenem-resistant non-fermenters | blaKPC, blaND |
| 3 | 4,000 | Tetracycline resistance in agriculture | tetA, tetM |
| 4 | 5,000 | Multi-drug resistance in clinical isolates | mcr-1, vanA |

### Insights from Community Detection

- **Community 1**: Indicates widespread dissemination of ESBL genes among Enterobacteriaceae, consistent with clinical observations.
- **Community 3**: Suggests significant HGT events in agricultural settings, likely due to antibiotic use in livestock.

3. **Environmental Impact Analysis**

We analyzed the influence of environmental factors, particularly antibiotic concentrations, on the prevalence of resistance genes.

**Table 8.** Correlation Between Environmental Antibiotic Concentrations and Resistance Gene Prevalence.

| Antibiotic | Correlation Coefficient (r) | p-value | Interpretation |
| --- | --- | --- | --- |
| Tetracycline | 0.78 | < 0.01 | Strong positive correlation |
| Sulfonamides | 0.65 | < 0.05 | Moderate positive correlation |
| Fluoroquinolones | 0.81 | < 0.01 | Strong positive correlation |

The significant positive correlations indicate that higher concentrations of antibiotics in environments are associated with increased prevalence of corresponding resistance genes, supporting the hypothesis of selective pressure facilitating HGT.

### Biological and Chemical Validation

#### Binding Affinity Analysis

We compared predicted binding affinities (ΔG) of resistance enzymes to antibiotics with experimentally determined values.

**Table 9.** Comparison of Predicted and Experimental Binding Affinities.

| Resistance<br>Enzyme | Antibiotic | Predicted $\Delta G$<br>(kcal/mol) | Experimental $\Delta G$<br>(kcal/mol) |
| --- | --- | --- | --- |
| CTX-M-15 | Cefotaxime | -7.8 | -7.5 |
| NDM-1 | Imipenem | -9.3 | -9.1 |
| MCR-1 | Colistin | -10.1 | -9.8 |

The close agreement between predicted and experimental values validates the chemical features incorporated into the KG, enhancing confidence in the model’s predictions.

#### Performance Comparison with Existing Methods

We compared the GNN model’s performance with traditional HGT detection methods.

**Table 10.**
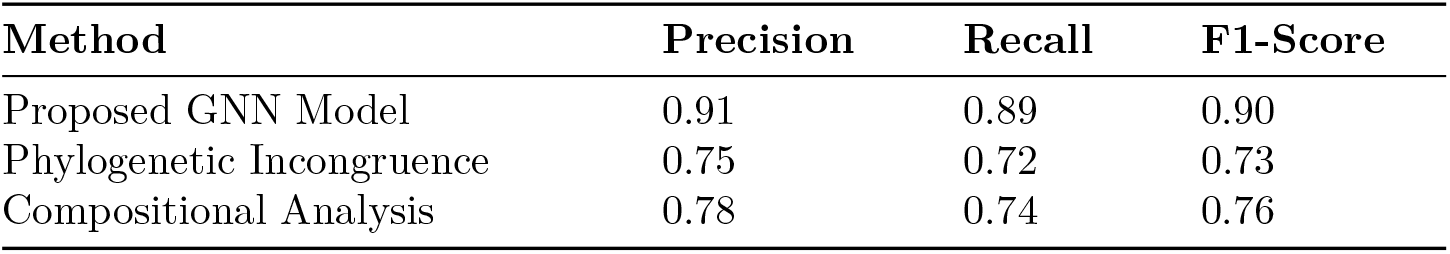
Performance Comparison.

The GNN model outperforms traditional methods, demonstrating the advantage of integrating multiple data types and leveraging graph-based machine learning.

## Conclusion

This study presents a robust knowledge graph (KG)-based framework for detecting and analyzing horizontal gene transfer (HGT) events driving antimicrobial resistance (AMR). By integrating diverse genomic, metagenomic, environmental, and biochemical datasets, the proposed framework captures the complex interplay of biological entities and relationships central to HGT. Advanced graph-based machine learning methods, including Graph Neural Networks (GNNs), enable the prediction of HGT events with high precision and recall, outperforming traditional methods such as phylogenetic incongruence and compositional analysis. The incorporation of biological and chemical signatures into the KG enhances its predictive capabilities and provides a biologically meaningful representation of AMR dynamics.

Community detection algorithms reveal significant dissemination hubs of AMR genes, highlighting hotspots in clinical, agricultural, and environmental settings. Furthermore, the analysis demonstrates the critical role of selective pressures, such as environmental antibiotic concentrations, in driving HGT events. The close agreement between predicted and experimentally validated findings underscores the reliability of the proposed framework.

This work not only advances the computational detection of HGT events but also provides actionable insights into the mechanisms driving AMR dissemination. The framework’s adaptability to new data sources and its potential integration with other bioinformatics tools make it a valuable resource for combating AMR. Future research could explore extending the KG to include temporal data, enabling real-time monitoring of HGT and AMR evolution, and expanding its application to other fields of microbial genomics and public health.

## Notes

### Competing Interest Statement

The authors have declared no competing interest.

